# Valine and Inflammation Drive Epilepsy in a Mouse Model of ECHS1 Deficiency

**DOI:** 10.1101/2024.06.13.598697

**Authors:** Meghan M. Eller, Aamir R. Zuberi, Xiaorong Fu, Shawn C. Burgess, Cathleen M. Lutz, Rachel M. Bailey

## Abstract

ECHS1 Deficiency (ECHS1D) is a rare and devastating pediatric disease that currently has no defined treatments. This disorder results from missense loss-of-function mutations in the *ECHS1* gene that result in severe developmental delays, encephalopathy, hypotonia, and early death. ECHS1 enzymatic activity is necessary for the beta-oxidation of fatty acids and the oxidation of branched-chain amino acids within the inner mitochondrial matrix. The pathogenesis of disease remains unknown, however it is hypothesized that disease is driven by an accumulation of toxic metabolites from impaired valine oxidation.

To expand our knowledge on disease mechanisms, a novel mouse model of ECHS1D was generated that possesses a disease-associated knock-in (KI) allele and a knock-out (KO) allele. To investigate the behavioral phenotype, a battery of testing was performed at multiple time points, which included assessments of learning, motor function, endurance, sensory responses, and anxiety. Neurological abnormalities were assessed using wireless telemetry EEG recordings, pentylenetetrazol (PTZ) seizure induction, and immunohistochemistry. Metabolic perturbations were measured within the liver, serum, and brain using mass spectrometry and magnetic resonance spectroscopy. To test disease mechanisms, mice were subjected to disease pathway stressors and then survival, body weight gain, and epilepsy were assessed.

Mice containing KI/KI or KI/KO alleles were viable with normal development and survival, and the presence of KI and KO alleles resulted in a significant reduction in ECHS1 protein. ECHS1D mice displayed reduced exercise capacity and pain sensation. EEG analysis revealed increased slow wave power that was associated with perturbations in sleep. ECHS1D mice had significantly increased epileptiform EEG discharges, and were sensitive to seizure induction, which resulted in death of 60% of ECHS1D mice. Under basal conditions, brain structure was grossly normal, although histological analysis revealed increased microglial activation in aged ECHS1D mice. Increased dietary valine only affected ECHS1D mice, which significantly exacerbated seizure susceptibility and resulted in death. Lastly, acute inflammatory challenge drove regression and early lethality in ECHS1D mice.

In conclusion, we developed a novel model of ECHS1D that may be used to further knowledge on disease mechanisms and to develop therapeutics. Our data suggests altered metabolic signaling and inflammation may contribute to epilepsy in ECHS1D, and these alterations may be attributed to impaired valine metabolism.

## Introduction

ECHS1 Deficiency (ECHS1D) is a rare, autosomal recessive disorder caused by loss-of-function mutations in *ECHS1*. Disease-associated variants influence various factors such as splice sites, mitochondrial localization, protein folding and stability, or substrate binding. Since its discovery in 2014, 40 variants have been reported.^1^ *ECHS1* is a nuclear DNA gene encoding enoyl-CoA hydratase, short chain 1 or ECHS1, which translocates to the inner mitochondrial matrix to catalyze the hydration of substrates for beta-oxidation of fatty acids (FAs) and breakdown of branched chain amino acids (BCAAs; valine, leucine and isoleucine).^2^

ECHS1D has a Leigh syndrome-like presentation including severe developmental delays, hypotonia, dystonia, epilepsy, and failure to thrive.^3^ The most common neurological features are basal ganglia lesions and increased brain lactate, although the severity is variable among individuals.^4^ As symptoms worsen, there is progressive volume loss in the cerebellum that extends throughout the brain.^4, 5^ Similarly, epileptic events become more frequent and severe over time.^6^ Histologically, neuroinflammation and vacuolization is commonly observed in mitochondrial disease patients and mouse models.^7, 8^ Only one ECHS1D patient has undergone post-mortem assessment to date^9^ and knowledge on ECHS1D neuropathology is limited.

ECHS1 has a multi-faceted role in metabolism,^2^ although patients typically only have increased valine-derived metabolites, suggesting a critical role of impaired valine breakdown in driving disease.^10^ Based on this, dietary reduction of valine intake has been performed in a few patients with variable success in symptom management.^10–14^ It is hypothesized that accumulation of toxic valine intermediates within mitochondria disrupt normal function.^15^ Measures of mitochondrial function in patient fibroblasts showed decreased activity of pyruvate dehydrogenase complex (PDC) and oxidative phosphorylation (OxPhos) complexes, but these findings are variable.^4, 15, 16^ Additional studies have demonstrated that ECHS1 knock-out (KO) cells have reduced OxPhos complex assembly and mitochondrial oxygen consumption, implicating respiratory defects in ECHS1D pathogenesis.^17^

In this study, we tested potential mechanisms underlying disease progression in a novel animal model of ECHS1D. Behavioral testing revealed that a significant reduction of ECHS1 in mice results in reduced exercise capacity and pain sensation. ECHS1D mice had disrupted EEG activity and sleep patterns, increased seizure susceptibility, and neuroinflammation. Interestingly, ECHS1D mice displayed normal energy status and OxPhos assembly in the liver, while amino acid abundance was significantly increased. Challenges with a valine-enriched diet or an acute inflammatory agent worsened the epileptic phenotype and induced death, implicating impaired valine oxidation and inflammation in disease pathogenesis.

These findings detail novel phenotypes of ECHS1D in mice and identify potential drivers of disease, the knowledge of which can aid in the development of therapeutic interventions.

## Materials and methods

### Experimental Animals

All procedures were performed in accordance with protocols approved by the Institutional Animal Care and Use Committee of the University of Texas Southwestern Medical Center (UT Southwestern) and the Jackson Laboratory, both AAALAC-accredited facilities. Targeted mutations in the ECHS1 gene were generated at the Jackson Laboratory on a C57BL/6J background using CRISPR/Cas9 genome editing. Briefly, single cell zygotes were electroporated with Cas9 protein, sgRNA, and a mutagenic donor oligonucleotide (Supplemental Table 1). Founder mice were generated, and genome editing scored after sequencing the targeted locus. Founders containing the F33S KI allele were mated to C57BL/6J mice to confirm germ line transmission in N1 progeny, and the resulting heterozygous KI mice were backcrossed once more to C57BL/6J. The viability of homozygous mice was evaluated by intercrossing N2 heterozygous mice. Two ECHS1 mutant strains were recovered and assigned with Jackson Laboratory Stock numbers. Stock 35254 (C57BL/6J-*Echs1^em1Lutzy^*/ Mmjax) contains the A31A and F33S edited alleles and Stock 35256 (C57BL/6J-*Echs1^em3Lutzy^*/ Mmjax) contains a single indel in exon 2 that is predicted to generate a frameshift null allele. Strain 35254 was homozygous viable with both male and female mice fertile. Strain 35256 was homozygous lethal (0 homozygous mice observed from 65 progeny generated from the intercross of heterozygous males and females). Breeders of both strains were transferred to UT Southwestern and maintained under controlled environmental conditions, 12-hour light-dark cycles, and provided food and water ad-libitum.

### ECHS1 Expression

At 2 months of age, mice were deeply anesthetized with tribromoethanol and perfused with PBS-heparin. Tissues were immediately frozen on dry ice and stored at –80°C. Samples were homogenized in 0.5% NP-40 buffer with protease inhibitor cocktail (Sigma, #11836170001), incubated on ice for 10 min, sonicated, and centrifuged at 12,000xrpm for 15 min at 4°C. Supernatant was stored at –80°C. Protein was quantified using Pierce BCA Protein Assay kit (Thermo Scientific, #23225). For western blot analysis, 45ug of protein was run on a 4-20% gel (BioRad, #4561096), transferred to a PVDF membrane and blocked with 5% milk in TBS-T. Blots were incubated with mouse anti-ECHS1 (1:1000; Proteintech, #89017583) followed by mouse peroxidase secondary antibody (1:10,000; Jackson Immuno #115-035-146). Protein loading was normalized to Actin (primary 1:10,000, Cell Signaling #4970S; secondary 1:10,000 Jackson Immuno #111-035-144) or GAPDH (primary 1:10,000, Meridian #H68504M; secondary 1:10,000 Jackson Immuno #115-035-146). Quantification was performed using ImageJ.

### Behavioral Testing

#### Rotarod

Mice are placed on a stationary rotarod (Columbus Instruments) which is then accelerated from 4 to 40 rpm over five min. The time that each mouse falls from the rod is recorded. If a mouse holds onto the rod and rotates completely around, it is recorded as a fall at that time. Each mouse is tested four times a day for two consecutive days with ∼30 min between trials.

#### Hot plate

Mice were placed onto a hotplate (IITC Life Science Inc.) with a surface temperature of 52°C. The latency until the mouse licked or shook the hind paw or jumped was recorded. The mouse was immediately removed from the hotplate once it had shown a response. The test was repeated with ∼30 min between trials.

#### Open Field

Mice were placed in the periphery of a novel open field (44 cm x 44 cm, walls 30 cm high) in a dimly lit room (∼60 lux) and allowed to explore for 10 min. Animals were monitored by a video camera and data analyzed via Ethovision software (13.0, Noldus) to determine the time, distance moved and number of entries into two areas: the periphery (5 cm from the walls) and the center (14 cm x 14cm).

#### Elevated Plus Maze

Mice were placed in the center of an elevated plus maze (each arm 30 cm long and 5 cm wide with two opposite arms closed by 25 cm high walls) elevated 31 cm in a dimly lit room (∼60 lux) and allowed to explore for 5 min. Animals were monitored by a video camera and data analyzed via Ethovision software (13.0, Noldus).

#### Fear Conditioning

Fear conditioning was measured in boxes equipped with a metal grid floor connected to a scrambled shock generator (Med Associates Inc). For training, mice were placed in the chamber for 2 min, then received 3 tone-shock pairings (30 sec white noise, 80 dB tone co-terminated with a 2 sec, 0.5 mA footshock, 1 min intertrial interval). The following day, context memory was measured by placing the mice into the same chambers for 5 min with freezing behavior automatically scored by the software. 48 hours after training, memory for the white noise cue was measured by placing the mice in an altered box (plastic floor, “V” ceiling, and a vanilla odor). Freezing in the novel context was measured for 3 min, the noise cue was turned on for an additional 3 min and freezing was measured.

#### Pre-Pulse Inhibition

Startle was measured using a SR-Lab Startle Response System (San Diego Instruments). Mice were placed into Plexiglas holders and allowed to acclimate to the chamber and background white noise (70 dB) for 5 min. After acclimation, startle stimuli (120 dB, 40 ms, white noise) were presented with an average interstimulus interval of 20 sec (range 13 - 27 sec). A prepulse stimulus (20 msec tone, 0, 4, 8 or 12 dB above the background noise) was presented 80 msec before the startle stimulus, in a pseudorandom order.

#### Endurance

All mice were familiarized to the treadmills for 2 days prior to testing. The acclimation paradigm was as follows: Day 1 - 5 min rest; 5 min, 8 m/min; 5 min, 10 m/min; Day 2 - 5 min rest; 5 min, 10 m/min; 5 min, 12 m/min. On Day 3, mice were placed on the treadmill for 5 min at rest, followed by running at 10 m/min for 40 min. Speeds were then increased at the rate of 1 m/min every 10 min until the speed reached 13 m/min, and finally increased at the rate of 1 m/min every 5 min until exhaustion. The exhaustion time was noted as the time at which the mice remained on the electric shock grid for 5 continuous seconds, without attempting to resume running.

#### PTZ Seizure Induction

The pentylenetetrazol (PTZ) kindling model was performed as described.^18^ Briefly, mice received an intraperitoneal (i.p.) injection of 30mg/kg PTZ dissolved in sterile saline every other day for a total of 12 injections. Mice were observed for 30 minutes following injection and seizure severity was scored by a blind observer using a standard Racine scale.

### Telemetry Recording

Mice were implanted with HD-X02 telemetry probes using intended coordinates LH: AP +1.0, ML -1.5, RH: AP -2.0, ML +2.0 (Data Systems International, DSI). After at least one week of recovery, dural recordings were acquired over a 24-hour period consisting of 1 dark and 1 light cycle. Data were acquired using Ponemah and analyzed in Neuroscore. Sleep scoring was performed using the Rodent Sleep Scoring 2 program and total power was automatically determined by the software. Epileptic spikes and spike trains were quantified using the Spike Analysis program.

### Histology

At 18 months of age, mice were deeply anesthetized with tribromoethanol and perfused with PBS-heparin. Whole brain was drop-fixed in 10% NBF for 48 hours then processed, paraffin embedded, and microtome sectioned at 5-micron thickness. IHC staining was performed using standard techniques. Tris or citric acid antigen masking solution (Vector Laboraties, #H3300, #H-3301) was used for antigen retrieval. Primary antibodies used were anti-GFAP (1:750; Abcam,

#ab53554) or anti-IbaI (1:1000; Abcam, #ab28319). Secondary antibodies were biotinylated anti-chicken or anti-goat secondary antibody (1:1000; Vector Labs, BA-1000). Staining was visualized with a Vecta stain ABC HRP Kit (Vector Lab, PK-6100) and DAB kit (Vector Labs, NC9276270). Counter staining with hematoxylin was performed prior to coverslipping with aqua polymount (Polyscience Inc., 18606-20). Hematoxylin and eosin staining was performed by the UT Southwestern Histopathology Core.

### Magnetic Resonance Spectroscopy

At 9 months of age, T2 FLAIR images were obtained using ^1^H frequency (300 MHz) on the Agilent (Varian) 7.0 T MR imaging system. A 3mm x 3mm x 3mm voxel was placed over the midbrain for spectroscopy measures. All metabolite concentrations were normalized to water.

### Liver Collection for Metabolite Measurements

Liver samples were collected from 3-month-old mice anesthetized using isofluorane. The liver was exposed without puncturing the diaphragm, then cut by the vessels and snap frozen in liquid nitrogen within 10 seconds of harvest.

### Measurement of Nucleotides and Short Chain Acyl-CoAs

Ion-pairing reverse-phase liquid chromatography ionization-tandem mass spectrometric was used to quantify adenine nucleotides and short-chain acyl-CoAs as previously reported.^19^ Frozen liver samples were spiked with stable-isotope-labeled ATP, AMP, acetyl-CoA, and malonyl-CoA internal standards and homogenized in 0.4 M HClO_4_ (PCA) containing 0.5 mM EGTA. Neutralized supernatants were subjected to LC-MS/MS analysis using a Shimadzu LC-20AD liquid chromatography (LC) system coupled to a Triple Quad^TM^ 5500+ QTrap LC-MS/MS mass spectrometer (Applied Biosystems/Sciex Instruments). A reverse-phase C18 column (Waters Atlantis T3, 150 x 2.1 mm, 3 mm) was used with an LC mobile phase consisting of water/methanol (95:5, v/v) with 4 mM DBAA (eluent A) and water/acetonitrile (25:75, v/v; eluent B). Multiple-reaction monitoring (MRM) in positive ion mode was used to detect each analyte and its stable isotope internal standard.

### Measurement of Amino Acids

Frozen liver samples were spiked with stable isotope labeled amino acid internal standards (Isotec) and quantified using LC-MS/MS as we previously reported.^20^ Tissue was homogenized in 4% perchloric acid, and supernatant was neutralized with 0.5 M K_2_CO_3_. After salt removal, supernatants were derivatized using established protocols.^21^ Amino acids were separated on a reverse phase C18 column (Waters, Xbridge, 150×2.1 mm, 3.0μm) with gradient elution and detected using the MRM mode by monitoring specific transitions under positive electrospray on a Triple Quad^TM^ 5500+ QTrap LC-MS/MS mass spectrometer (Applied Biosystems/Sciex Instruments). Amino acids were quantified by comparing individual ion peak areas to that of the stable isotope internal standard.

### OxPhos Complex Assembly

Mitochondria were isolated from 12-month-old mouse liver. Tissues were homogenized in isolation buffer containing protease inhibitor (5mM HEPES, 70mM sucrose, 220mM mannitol, 5mM MgCl2, 10mM KH2PO4, 1mM EGTA; pH 7.2) and centrifuged as previously described.^22^ Protein content was measured with the Bradford assay. 50ug of protein was solubilized in sample buffer (Invitrogen BN2003) containing 5% digitonin (Sigma D141) and centrifuged at 15,000xg for 10 minutes. G-250 coomassie additive (Invitrogen BN2004) was added to the supernatant. Protein was loaded on a 3-12% gel (Invitrogen BN10001B0X) and run using Invitrogen buffers (BN2002, BN2001) as described (Ref: Nishihara).^23^ The gel was stained with Coomassie to image the NativeMark ladder (Thermofisher, #LC0275), then destained and transferred onto a PVDF membrane. The membrane was fixed^24^ and blocked in 5% BSA in TBST, incubated in OxPhos cocktail (1:5000; Abcam, ab110413) and then mouse peroxidase secondary antibody (Jackson Immuno #115-035-146) in 5% BSA in TBST. Protein loading was normalized to the Complex II band.

### Valine Diet

A custom diet containing 3.8% valine was formulated by Envigo using their standard 2016 diet as a base and stored at 4°C. Concentrations of other chow components were the same. Valine chow was provided to breeders and their weaned litters.

### Lipopolysaccharide Challenge

Lipopolysaccharide from Escherichia coli O55:B5 (Sigma, L2880) was dissolved in sterile PBS at 0.5mg/mL. At 2 months of age, baseline weights were recorded, and mice received an i.p. injection of 5mg/kg LPS. Weights were collected daily for the first week following injection, and once a week thereafter.

### Imaging and Analysis

Light microscopy images were captured at 20X on a Hamamatsu Nanozoomer 2.0 HT. Percent GFAP staining was quantified using the area quantification algorithm in HALO Imaging Analysis Platform (Halo2.2. Indica Labs, Albuquerque, NM, USA). Microglia counts were determined using the microglial activation algorithm within HALO. The striatum was outlined for analysis and the same outline and threshold settings were applied to each image in the respective groups.

### Statistics

One-way or two-way ANOVA with Tukey or Sidak’s multiple comparison was used for multiple group comparisons and Student’s t-test was used for comparisons between two groups. Geisser-Greenhouse’s epsilon correction was used if the estimated epsilon was < 0.75. Survival differences were analyzed using the log rank test. Statistical significance was set at p ≤ 0.05. Outliers were determined using Grubbs’ test. Statistical analysis and graphing were done using GraphPad Prism software (v9.5.0; GraphPad Software). Values are expressed as mean +/- SEM.

## Results

### Development of ECHS1D Mouse Model

#### Generation of ECHS1D Mice

There are currently no published animal models of ECHS1D and complete ECHS1 knockout is embryonic lethal. While heterozygous knockout mice possess a 50% reduction in ECHS1 expression, they develop normally with only mild lipid accumulation and a cardiomyopathy phenotype.^25, 26^ To establish a model of disease, CRISPR/Cas9 was used to generate a knock-in (KI) mouse line that contains a disease-associated variant (c.98T>C; p.F33S) (Fig. 1).^4^ A knock-out (KO) line was also generated that possesses a frameshift mutation in exon 2 that results in an early stop codon. Heterozygous KI mice were bred together to generate homozygous KI mice, or with heterozygous KO mice to establish the compound heterozygous ECHS1D mouse as depicted in Fig. 1. While we were unable to obtain homozygous KO mice due to embryonic lethality, KI/KI and KI/KO mice were viable and fertile. Sanger sequencing was used to confirm germline mutations in mice (Fig. 1 B-E).

**Figure 1:**
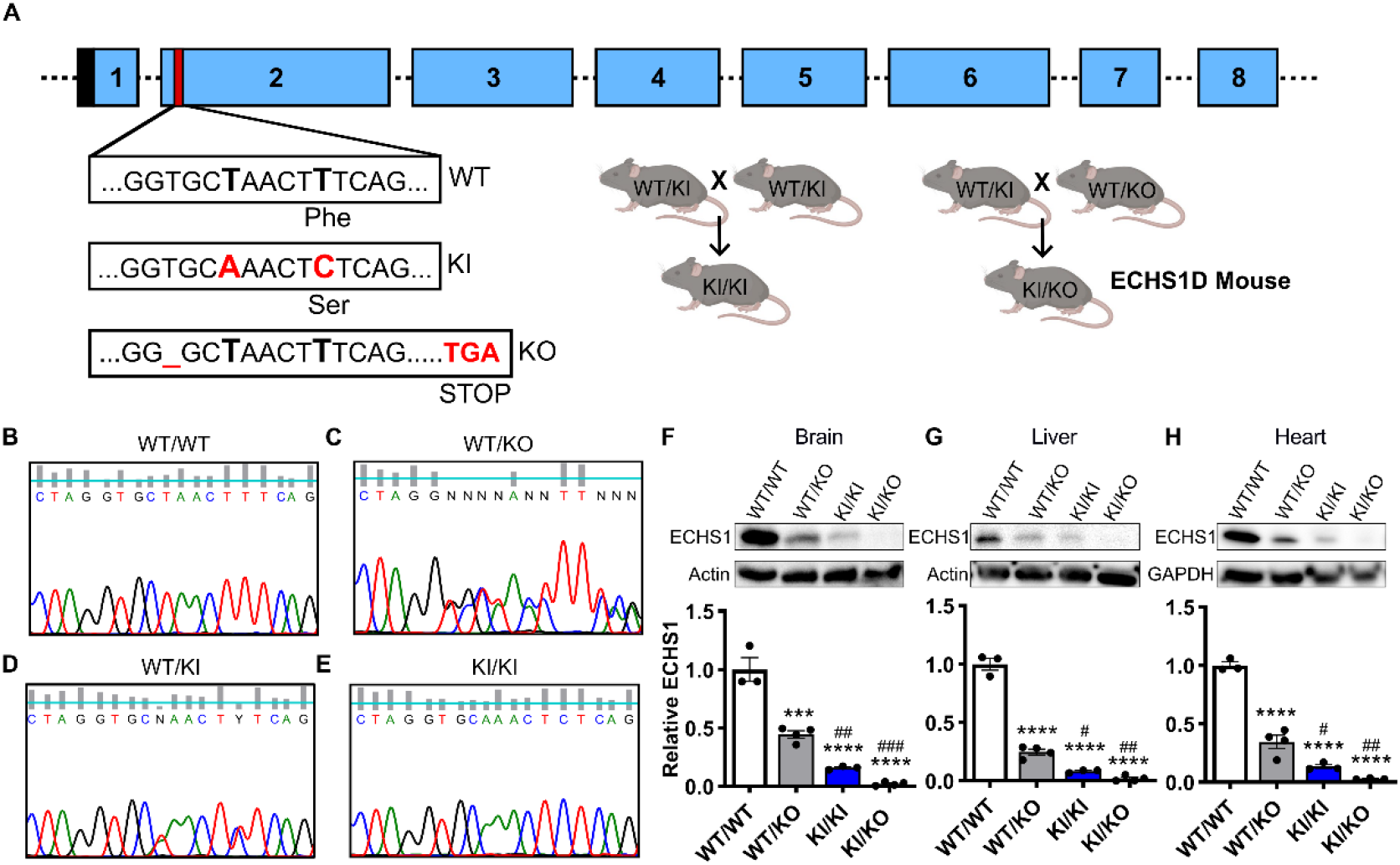
Generation and validation of ECHS1 Deficiency mouse model. ECHS1D mice were generated from *Echs1* knock-in (KI) and *Echs1* knock-out (KO) lines. (**A**) Depiction of mouse *Echs1* gene, with exons 1-8. Black = mitochondrial localization signal; red = target region. The F33S KI allele has a synonymous mutation (c.93T>A; p.A31A) and a missense mutation (c.98T>C; p.F33S). The KO allele has a frameshift mutation (c.90delT; p.G30del). Heterozygous KI mice (WT/KI) were bred to establish homozygous KI mice (KI/KI). WT/KI mice were also bred with heterozygous KO (WT/KO) mice to generate the KI/KO line. (**B-E**) Sequencing of genomic DNA showing the presence of the mutations within transgenic mice. ECHS1 expression in tissues was assessed in 2-month-old mice (N=3-4/genotype). (**F-H**) Total protein from brain (F), liver (G), and heart (H) was analyzed via western blot. Band intensity was quantified and ECHS1 was normalized to actin or GAPDH and set relative to WT levels. Each dot represents an individual mouse with SEM indicated. One-way ANOVA with Tukey’s post-hoc analysis, ***p<0.001, ****p<0.0001 compared to WT/WT; ^#^p<0.05, ^##^p<0.01, ^###^p<0.001 compared to WT/KO.

#### Model Validation

In ECHS1D patients, genetic variants result in the loss of ECHS1 function.^16^ To determine if the introduced mutations affect ECHS1 expression in our model, protein was quantified in brain, liver, and heart (Fig. 1). The KI mutation significantly decreased ECHS1 protein in a gene-dose dependent fashion (Fig. 1F-H). Across tissues, WT/KO mice retained 50% of WT ECHS1 as expected, while KI/KI mice expressed ∼12% and KI/KO mice had 3% of WT levels. As KI/KI mice were largely phenotypically normal and with higher ECHS1 expression, subsequent assessments focused on KI/KO mice, now referred to as ECHS1D mice.

### Behavioral Deficits in ECHS1D Mice

#### Motor Coordination and Memory

ECHS1D mice developed normally, without an outward phenotype, and had similar body weight as WT littermates (Supplemental Fig 1). Mice were behaviorally tested to assess motor, sensory, and cognitive function starting at 3 months of age (Fig. 2, Supplemental Fig. 2). Rotarod testing was used to evaluate motor learning and coordination, as motor skills are commonly impaired in ECHS1D patients.^3^ In the first trial, performance was similar between groups demonstrating intact motor function in ECHS1D mice. ECHS1D and WT mice had similar improvements in performance, supporting normal motor learning in this model (Supplemental Fig. 2A). With repeated testing, however, ECHS1D mice progressed to having a significantly reduced latency to fall on trial 8 (Fig. 2A). In open field and elevated plus maze testing, all mice traveled similar distances and spent similar time in each testing area, suggesting that impaired rotarod performance was unlikely due to differences in overall movement or anxiety (Supplemental Fig 2B-E). During fear conditioning, there were no differences in percent freezing during the context or cue portion, indicating that short term memory was typical in ECHS1D mice (Supplemental Fig 2F-G). In longitudinal testing deficits generally did not worsen, suggesting that under normal conditions behavioral phenotypes in ECHS1D mice are not progressive (Supplemental Fig. 3). The results at later time points may have been confounded by learning from repeat testing, so future studies should test naïve animals at each age.

**Figure 2:**
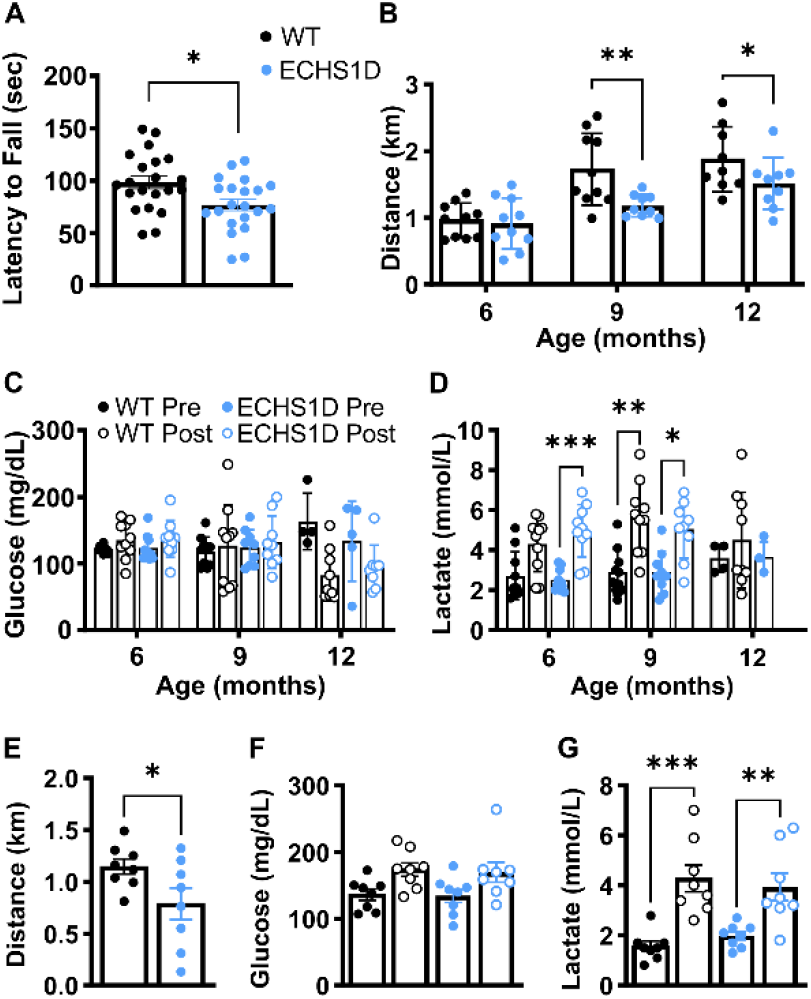
Exercise capacity is reduced in ECHS1D mice. **(A)** At 3 months of age, WT and ECHS1D mice (N=20/genotype) were subjected rotarod testing and latency to fall is shown for trial 8. (**B**) A separate cohort of WT and ECHS1D mice (N=10/genotype) were subjected to the treadmill endurance test to measure exercise capacity with repeat testing at 6, 9 and 12 months of age. The total distance run on an accelerating treadmill before exhaustion was recorded. (**C, D**) Blood glucose and lactate levels were measured before and after endurance testing at each test age. (**E-F**) Endurance testing was performed in a naïve cohort of mice (N=8/genotype) at 9 months of age. Total distance (E), blood glucose (F), and blood lactate (G) were measured. Each dot represents an individual mouse with SEM indicated. A, E: Student’s t-test. B-D, F-G: Repeat measure two-way ANOVA with Sidak multiple comparisons. *p<0.05, **p<0.01, ***p<0.001.

### Sensory Responses

Peripheral neuropathy is common among mitochondrial disease patients, which manifests as hyperalgesia or reduced sensation, often from demyelination.^27–29^ To measure nociception in our model, hotplate testing was performed. Latency to withdraw was comparable between genotypes in the first trial, but was significantly increased in ECHS1D mice compared to WT by the third trial, suggesting abnormal nociception (Supplemental Fig 2H). This alteration in sensory processing was specific to noxious heat, as there were no differences in sensory gating during pre-pulse inhibition using auditory stimuli (Supplemental Fig 2I). ECHS1D mice had increased withdraw latency in hotplate testing again at 12 months, which could indicate progression of the peripheral neuropathy (Supplemental Fig. 3I).

### Exercise Intolerance

Results from rotarod testing suggest exercise fatigue in ECHS1D mice, so we directly measured this with repeated endurance testing (Fig. 2B-D). While run distance was comparable between WT and ECHS1D mice at 6 months, ECHS1D mice ran significantly shorter distances at 9 and 12 months of age, supporting that ECHS1D mice have exercise intolerance (Fig. 2B). Blood glucose and lactate were measured before and after exercise, except for a subset of 12-month-old mice where the baseline measurement was lost due to sampling error. ECHS1D mice had similar pre- and post-exercise glucose levels as WT mice, indicating normal gluconeogenesis (Fig. 2C). ECHS1D and WT mice had similar pre-exercise lactate concentrations, which then increased following exercise (Fig. 2D). In WT mice, lactate was only significantly increased after exercise at 9 months, while ECHS1D lactate levels were significantly increased at both 6 and 9 months of age (Fig. 2D).

To determine if exercise intolerance in ECHS1D mice resulted from repeat testing, a group of naïve 9-month-old mice were also tested. In agreement with the longitudinal cohort, ECHS1D mice had significantly reduced run distances compared to WT mice, demonstrating that exercise intolerance at 9 months of age is present regardless of repeat testing (Fig. 2E). Additionally, blood glucose and lactate levels in the naïve cohort matched those in the longitudinal exercise cohort (Fig. 2F-G).

### Neurological Phenotype in ECHS1D Mice

#### EEG Activity

EEG recordings of Leigh syndrome patients show disorganized background neural activity,^30^ and ∼40% of ECHS1D patients develop epilepsy throughout their disease progression.^3^ To assess brain activity of ECHS1D mice, we performed EEG and EMG for 24 hours using wireless telemetry implants. Overall, ECHS1D mice had significantly elevated delta and theta power compared to WT mice (Fig. 3A). When examined within sleep stages, power was increased in wake and active wake only (Supplemental Fig. 4). To determine if these power alterations affect sleep, the percent time spent in each sleep stage was calculated. ECHS1D mice spent significantly more time in the active awake stage, where animals are awake and moving, and similar amounts of time in the wake stage, where animals are awake but not active, as compared to WT mice. Conversely, ECHS1D mice had a significantly reduced percentage of slow wave sleep compared to WT mice, although paradoxical sleep was similar (Fig. 3B).

**Figure 3:**
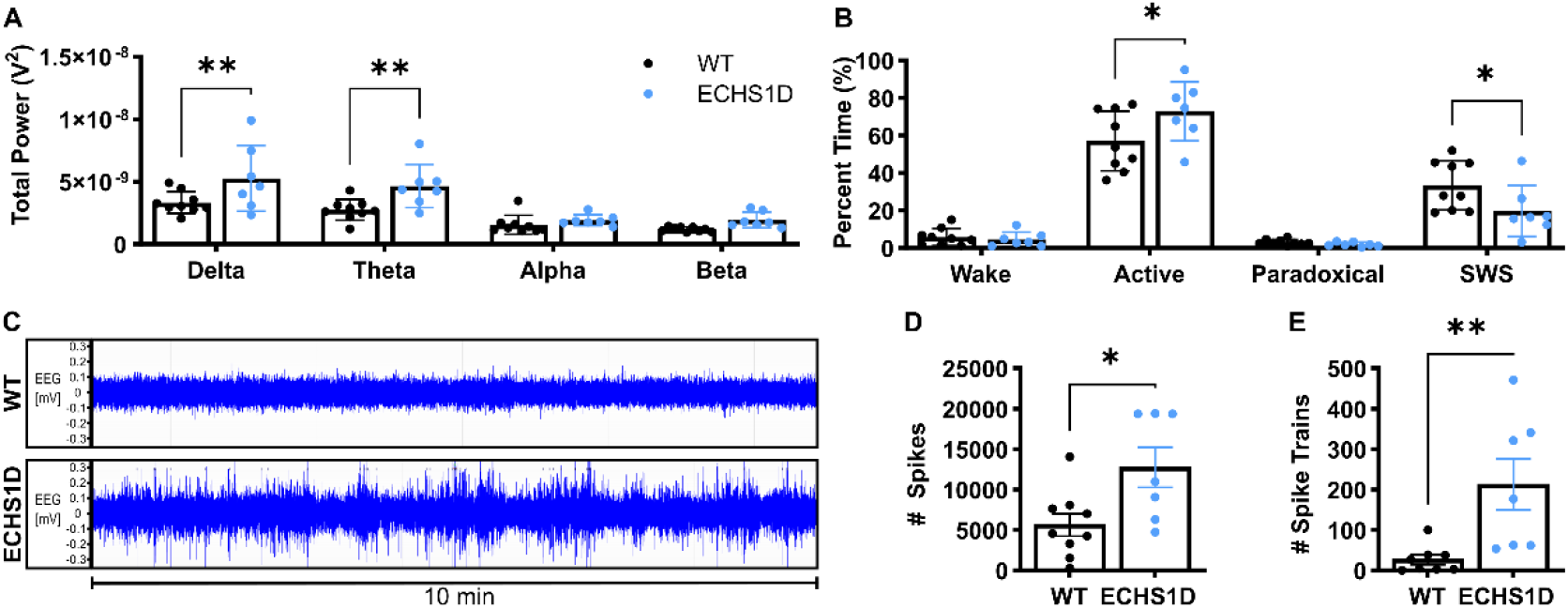
ECHS1D mice have abnormal baseline EEG. At ∼3.5 months of age, mice received wireless telemetry implants and EEG activity was recorded over a 24-hour period (N=7-9/genotype). (**A**) Total EEG power from delta, theta, alpha, or beta frequencies. (**B**) Sleep staging and percent time spent in wake, active wake, paradoxical sleep, or slow wave sleep was calculated. (**C**) Representative raw EEG traces from WT and ECHS1D. (**D-E**) Individual spikes (D) and spike trains (E) were quantified. Each dot represents an individual mouse with SEM indicated. A-B: Two-way ANOVA with Sidak’s multiple comparisons. D-E: Student’s t-test. *p<0.05, **p<0.01.

Representative EEG traces shown in Figure 3C demonstrate that ECHS1D mice have significantly increased EEG signal compared to WT mice. Quantification of epileptic spikes and spike trains confirmed a significant increase in epileptiform discharges in ECHS1D mice as compared to WT (Fig. 3D-E).

#### Seizure Susceptibility

Spontaneous seizures were not observed in ECHS1D mice, however, the altered baseline EEG suggests ECHS1D mice may be more susceptible to seizure onset. To test this, we performed a kindling induction paradigm using pentylenetetrazol (PTZ).^18^ Seizure severity was scored using the Racine scale depicted in Figure 4A. With the low dose used, WT mice failed to develop seizures. In contrast, ECHS1D mice developed progressive seizures that worsened over time with a significant reduction in latency to seize (Fig. 4B-C). Severe seizures in ECHS1D were associated with early lethality where 60% of mice died by injection 12, while WT mice had 100% survival (Fig. 4D). These results demonstrate that under standard conditions ECHS1D mice have abnormal brain activity that is associated with increased seizure susceptibility following PTZ treatment.

**Figure 4:**
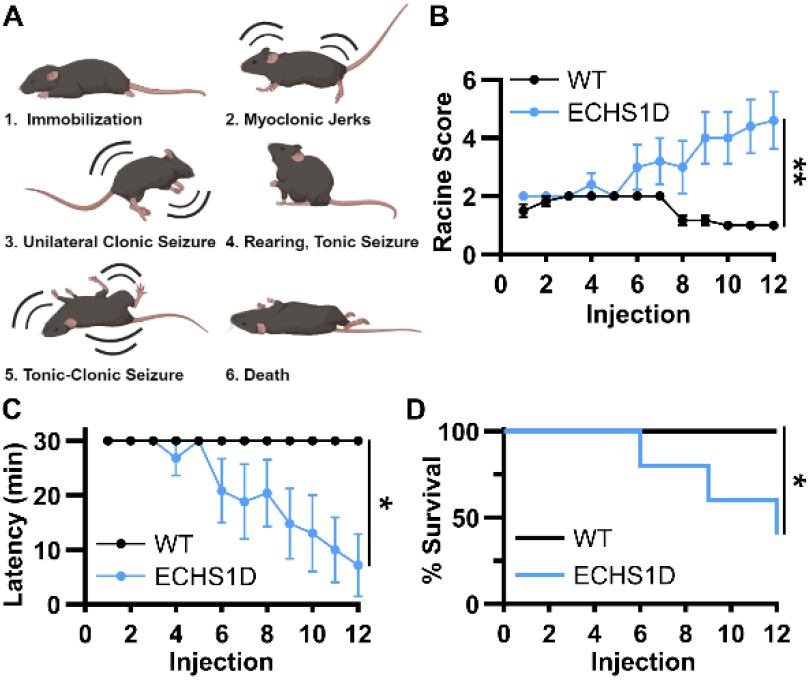
ECHS1D mice have increased seizure susceptibility. At ∼3.5 months of age, mice were subjected to pentylenetetrzaol (PTZ) seizure induction (N=5-6/genotype). (**A**) Depiction of Racine scale used to score seizure severity. (**B-D**) Mice received 12 injections of PTZ (30mg/kg) every other day and seizure severity (B), latency (C), and survival (D) was recorded following each injection. B-C: Two-way ANOVA, Genotype effect *p<0.05, **p<0.01. D: Log rank test, *p<0.05.

#### Brain Lactate Levels

ECHS1D patients often present with elevated circulating lactate^4, 31, 32^ and brain lactate.^15^ Although there were no differences in circulating lactate between ECHS1D and WT mice (Fig. 2, Supplemental Fig. 5), blood lactate does not reliably inform on brain lactate.^33^ To quantify brain lactate we performed magnetic resonance spectroscopy (MRS) on 9-month-old mice. While there was a trending increase of brain lactate in ECHS1D mice, this effect was variable (Supplemental Fig. 5B). We also found that concentrations of N-acetylaspartate (NAA), creatine, and choline were similar between groups, and the brain appeared structurally normal in ECHS1D mice (Supplemental Fig. 5C-D). Together this data suggests that lactic acidosis likely does not significantly contribute to the baseline ECHS1D mouse phenotype.

### Neuropathology in ECHS1D Mice

#### Brain Structure

To further assess the neurological phenotype in ECHS1D mice, brains were obtained from 18-month mice for histological analysis. This age was chosen as MRI scans at 9 months of age did not reveal gross morphological abnormalities (Supplemental Fig. 5) and basal ganglia degeneration is progressive in patients.^4^ Hematoxylin and eosin (H&E) staining confirmed that brain structures were similar between WT and ECHS1D mice, even at this late stage (Fig. 5A-B). Taken with our MRS findings, this supports that in the absence of stressors, ECHS1D mice have normal gross brain development and do not undergo overt neurodegeneration.

**Figure 5:**
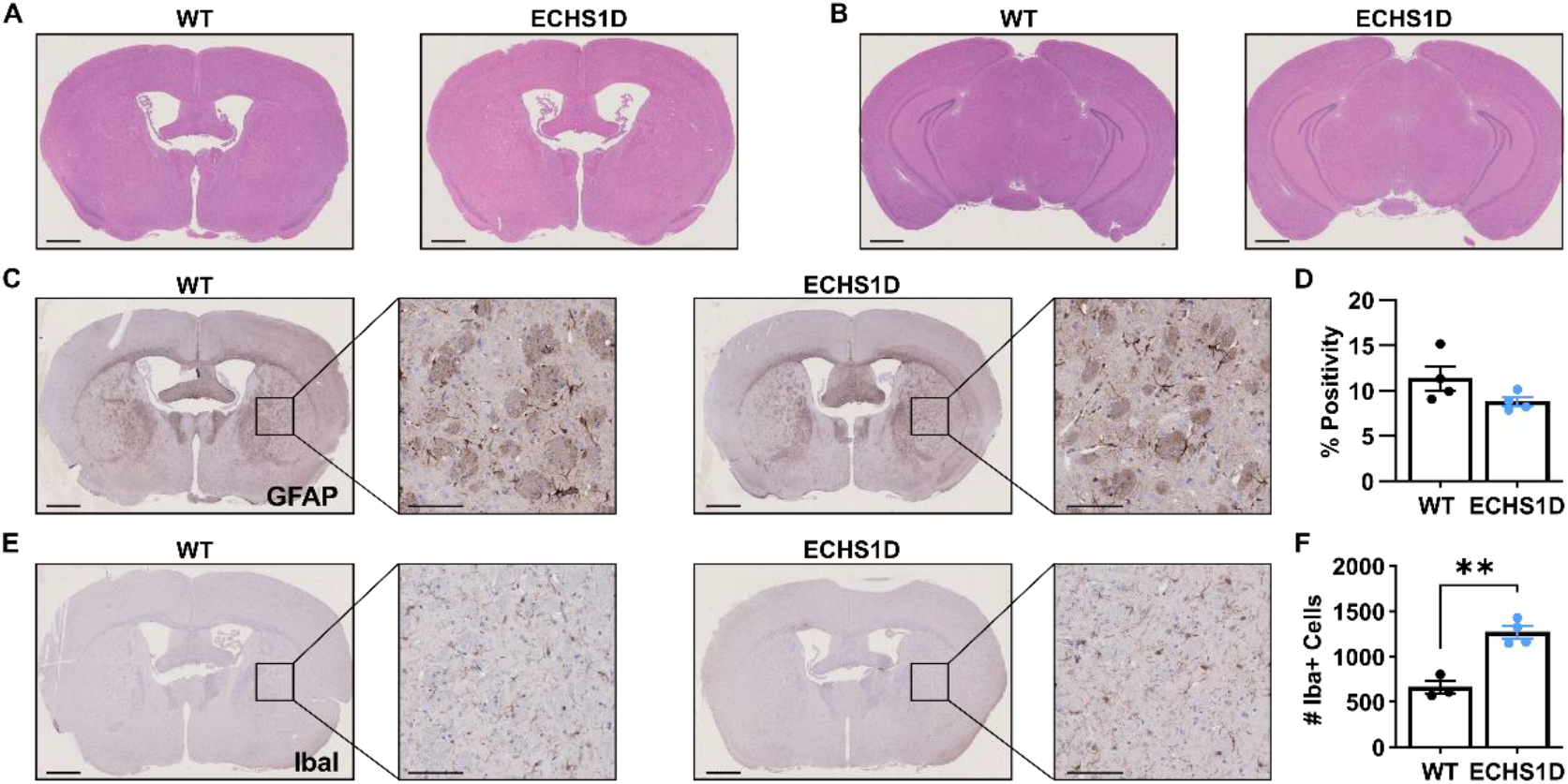
Aged ECHS1D mice have increased microglia. (**A, B**) Hematoxylin and eosin (H&E) staining was performed on WT and ECHS1D brain sections to assess overall brain structure (N=4/genotype; 18 months old). (**C, D**) GFAP staining was performed to assess astrocyte reactivity and percent positive staining within the striatum was quantified. (**E, F**) IbaI staining was performed to visualize microglia, and the number of microglia within the striatum was quantified. Scale bars = 1mm (whole brain images) and 100µm (insets). Student’s t-test, **p<0.01.

#### Neuroinflammation

Neuroinflammation is hypothesized to contribute to disease progression in Leigh syndrome and Leigh-syndrome like disorders,^34, 35^ but this has not been directly examined in ECHS1D patients. To determine if neuroinflammation is increased in ECHS1D mice, staining against GFAP and IbaI was performed to examine astrocyte and microglia reactivity, respectively. We quantified staining within the striatum and found a trending decrease in GFAP levels in ECHS1D mice (Fig. 5C-D). In contrast, IbaI staining was significantly increased within the striatum, where ECHS1D mice had twice as many microglia as WT mice (Fig. 5E-F). This data suggests that loss of ECHS1 may result in astrocyte death with a concurrent increase in microglial activation.

### Metabolic Abnormalities in ECHS1D Mice

#### Energy and Amino Acid Metabolism

As a nutrient-sensing enzyme,^36^ changes in ECHS1 activity can cause metabolic dysregulation.^37, 38^ To assess metabolism in ECHS1D mice under basal conditions, energy metabolites and amino acids were measured in tissues from 3-month-old mice (Fig. 6). Due to technical limitations in collecting brain tissue quickly enough to preserve labile metabolites,^19^ liver was assessed. Concentrations of acyl-CoA metabolites were unchanged between WT and ECHS1D mice, and energy charge and redox status were also similar (Fig. 6A). Due to the role of ECHS1 in BCAA catabolism, we also measured liver amino acid content. Levels of valine, isoleucine, and leucine were elevated in ECHS1D mice, but this was not significant. Interestingly, ∼60% of non-BCAA amino acids were significantly increased in ECHS1D mice compared to WT mice, suggesting widespread perturbations in overall amino acid metabolism (Fig. 6B). This data shows that, without ECHS1 activity, there is increased liver amino acid load, which may support maintenance of energy levels in ECHS1D mice.

**Figure 6:**
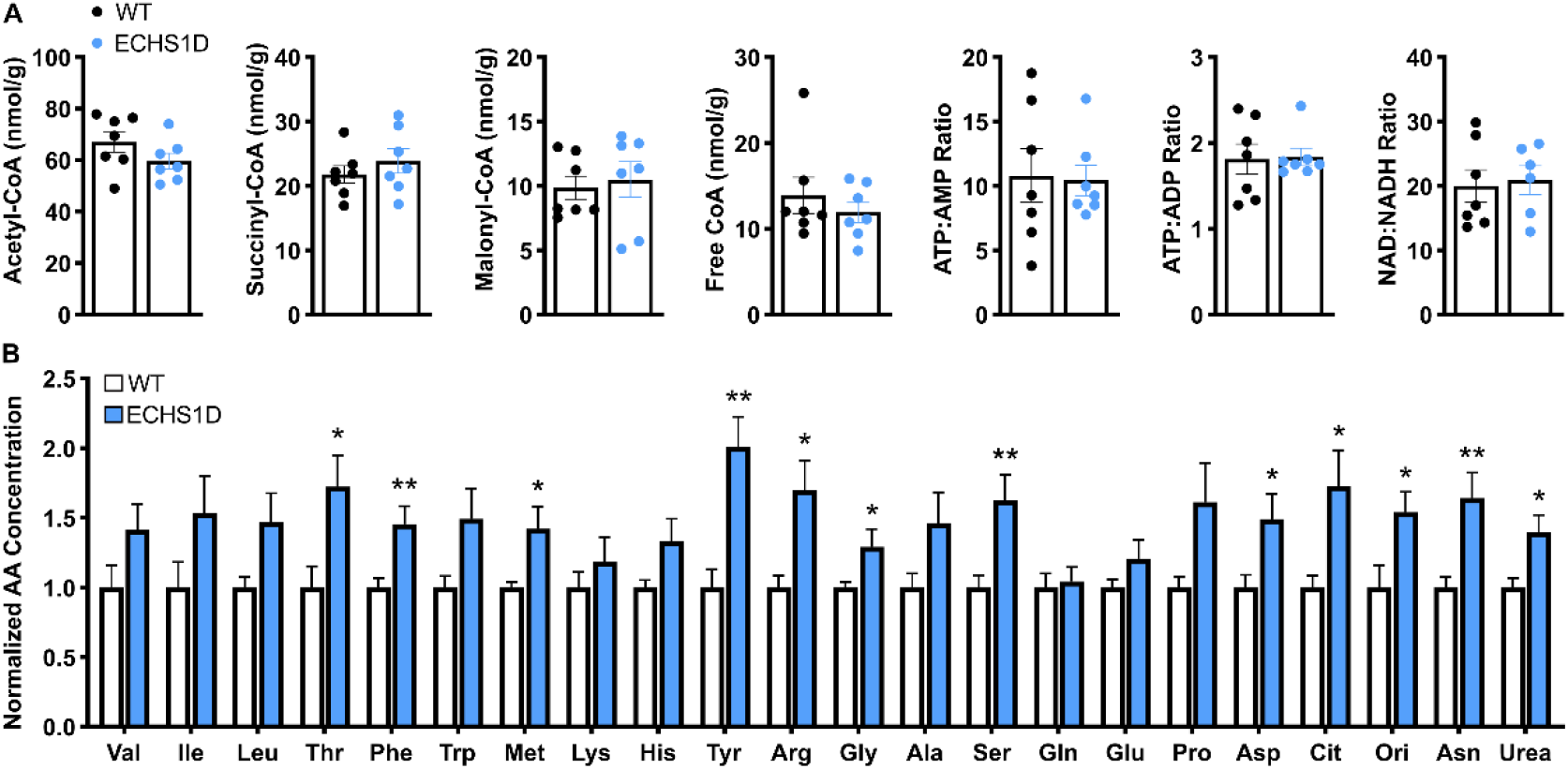
Liver amino acid concentrations are elevated in ECHS1D mice. Liver samples from 3-month-old WT and ECHS1D mice (N=7/genotype) were analyzed via GC-MS for quantification of (**A**) energy metabolites and amino acids (**B**). Amino acids were analyzed separately and graphed together; Student’s t-test, *p<0.05, **p<0.01.

#### Mitochondrial Complex Assembly

Recent findings showed that ECHS1-KO cells and ECHS1D patient fibroblasts have reduced OxPhos complex assembly.^17^ To test if this was recapitulated in ECHS1D mice, complex assembly was assessed in liver samples. There were no differences in the assembly of individual complexes or supercomplexes between WT and ECHS1D mice, suggesting normal OxPhos activity in the liver (Supplemental Fig. 6). This data suggests that under basal conditions, liver mitochondrial impairment likely does not contribute to disease presentation, although it remains unknown whether this is true for other tissues (e.g. muscle and brain) and under stress conditions.

### Disease Drivers

#### Dietary Valine

In ECHS1D, accumulation of reactive valine intermediates is thought to be toxic and dietary valine restriction is proposed to stabilize patient symptoms.^10–14^ To test if dietary valine supplementation exacerbates the disease phenotype of ECHS1D mice, a custom diet containing a 3% increase in valine content was formulated and provided to breeders (Fig. 7A). The resulting WT and ECHS1D pups were maintained on this diet throughout their life, and they had normal survival and body weight gains (Fig. 7B-C). As expected, serum valine was significantly increased in mice on the valine diet compared to mice on control diet, although there was no difference in serum valine levels of WT and ECHS1D mice on the same diet (Fig. 7D). To assess if dietary valine worsened the baseline epileptic phenotype, mice were subjected to PTZ seizure induction. In agreement with our prior PTZ experiment, ECHS1D mice on control diet developed progressively severe seizures that reduced survival by 50%, while WT mice were unaffected (Fig. 7E-G). Remarkably, valine treatment significantly exacerbated the seizure susceptibility of ECHS1D mice, who developed seizures following PTZ injection 1 that rapidly progressed in severity to result in 100% lethality by injection 9. This effect was specific to ECHS1D mice, as valine supplementation did not impact seizure susceptibility in WT mice, who remained seizure free (Fig. 7E-G). This data supports that increased dietary valine drives disease progression in ECHS1D.

**Figure 7:**
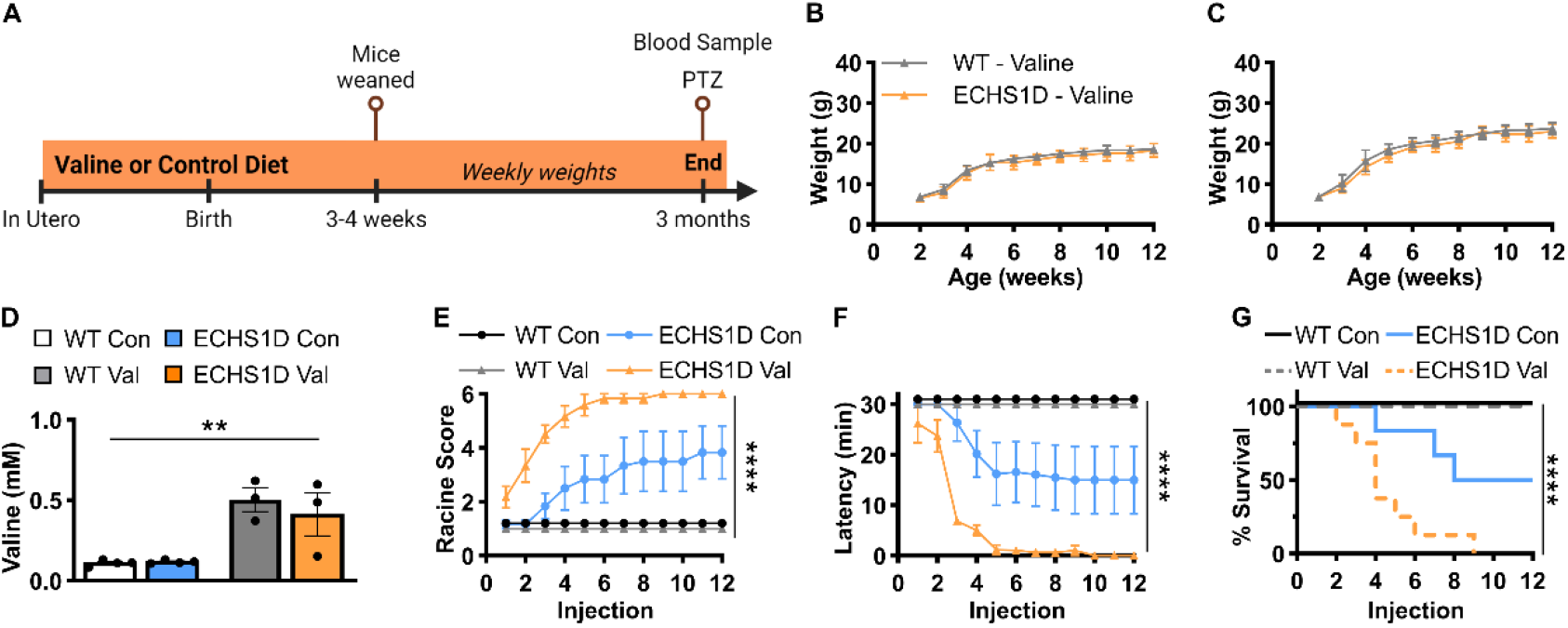
Dietary valine supplementation exacerbates seizure susceptibility in ECHS1D mice. Breeding WT/KI and WT/KO mice were given control (Con) or valine (Val) supplemented chow, and the resulting litters were maintained on the given diet throughout the study. (**A**) Study scheme. (**B, C**) Body weights of female (B) and male (C) mice. (**D**) Valine concentration within serum was quantified by GC-MS (N=3/group). (**E-G**) Mice (N=5-6/group) received 12 injections of PTZ (30mg/kg) every other day and seizure severity (B), latency to seize (C), and survival (D) was recorded following each injection. D: One-way ANOVA. E-F: Repeat measure two-way ANOVA with Tukey‘s multiple comparisons, Genotype effect. G: Log rank test. **p<0.01, ****p<0.0001.

#### Inflammatory Insults

The relationship between inflammation and mitochondrial dysfunction has been well documented,^39^ where mitochondrial disease patients often report a worsening of symptoms after infection, suggesting increased sensitivity to inflammatory stress.^3, 7^ To assess if inflammation worsens symptoms of ECHS1D mice, animals were given a single intraperitoneal injection of lipopolysaccharide (LPS). LPS causes an acute inflammatory response by triggering the release of inflammatory cytokines in various cell types and results in long-lasting neuroinflammation.^40^ Mice were monitored for weight loss and survival for 1-month following injection and then subjected to PTZ seizure induction (Fig 8A). Both control and ECHS1D mice had similar weight reductions 2 days after LPS treatment, but where control mice returned to baseline weight at 4 days post-injection, ECHS1D mice took 6 days to return to baseline (Fig. 8B). While LPS treatment was well-tolerated in control mice with 100% survival, 33% of ECHS1D mice died by 5 days post-injection (Fig 8C). Furthermore, the surviving ECHS1D mice exhibited failure to thrive in the weeks following injection where their body weights remained significantly decreased compared to control mice (Fig 8D). This data demonstrates that like patients, ECHS1D mice are hypersensitive to an acute inflammatory response that results in early lethality and regression.

**Figure 8:**
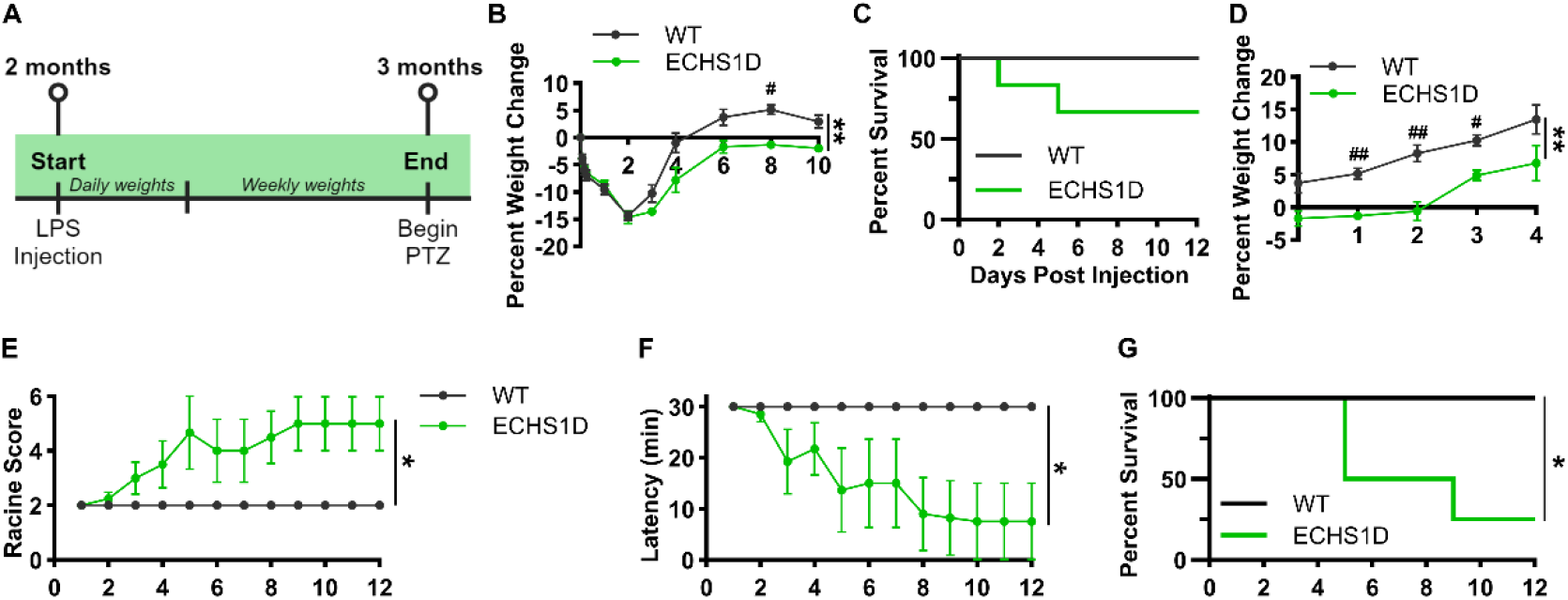
ECHS1D mice have failure to thrive and increased seizure susceptibility following LPS treatment. At 2 months of age, WT and ECHS1D mice received an intraperitoneal injection of lipopolysaccharide (LPS, 5 mg/kg; N=6/genotype). (**A**) Study design. (**B**) Percent change in body weight in the first 10 days following injection. (**C**) Percent survival following LPS treatment. Survival rates were stable past day 12. (**D**) Percent change in weekly body weight up to 1-month post-injection. (**E-G**) Surviving mice underwent seizure induction, and seizure severity (E), latency (F), and survival (G) was recorded following each injection. B, D-F: Repeat measure two-way ANOVA with Sidak’s multiple comparisons, Genotype effect *p<0.05, **p<0.01, WT vs. ECHS1D ^#^p<0.05, ^##^p<0.01. G: Log rank test, *p<0.01.

To determine if LPS treatment exacerbated the seizure susceptibility of ECHS1D mice, PTZ kindling was performed in surviving mice. WT mice did not develop any seizures, but LPS-treated ECHS1D mice had severe seizures (4 on Racine scale) by injection 3 that resulted in 25% survival (Fig 8E-G). Of note, in our prior experiments under basal conditions ECHS1D mice did not develop seizures until injection 6 and had 40-50% survival (Fig. 3, Fig. 8). While we cannot make a direct comparison across studies, this study suggests that acute neuroinflammation exacerbates seizure susceptibility in surviving ECHS1D mice.

## Discussion

This study is the first to report a mouse model for ECHS1 Deficiency, a rare Leigh-syndrome like disorder. The presence of the F33S knock-in variant significantly decreased ECHS1 protein levels which was compounded by the knock-out allele. To date, one patient has been reported to have the F33S variant along with the N59S variant.^4^ ECHS1 expression was significantly reduced in patient fibroblasts, similar to what we observed in our mice, although it is unknown how the N59S variant impacted ECHS1 expression. This patient presented with epileptic seizures within the first few days of life followed by developmental delays and hypotonia and was still reported to be alive at 3 years of age in 2015.^4^ Due to the lethality of *Echs1* knockout, ^25, 26^ the effects of ECHS1 reduction have only been minimally investigated in heterozygous knockout mice. A 50% reduction in ECHS1 causes cardiomyopathy phenotypes and lipid accumulation within liver and kidney.^25, 26^ Cardiac impairments are not observed in all patient reports and are mostly limited to severe neonatal cases.^3, 4, 15, 32^ Herein we focused on assessing neurological phenotypes as ECHS1D patients present with Leigh-like symptoms, however, future studies should assess cardiac function in ECHS1D mice.

ECHS1D mice had abnormal nociception that was specific to noxious heat. While peripheral neuropathy has not been reported in ECHS1D patients, it is estimated that one-third of mitochondrial disease patients develop neuropathy symptoms,^29^ and 81% of Leigh syndrome patients have abnormal peripheral nerve conduction.^27^ It is proposed that mitochondrial dysfunction within Schwann cells is responsible for peripheral demyelination.^41^ Based on our findings and supporting literature, we predict that ECHS1D patients and mice may present with peripheral demyelination and reduced nerve conduction velocity, though this needs to be tested.

Atypical EEG activity is common among mitochondrial disorders, with generalized slowing marked by increased low frequency power as the primary manifestation.^30, 42, 43^ ECHS1D mice had significantly elevated delta and theta power, specifically during wake and active wake sleep stages. Delta and theta power typically increase with extended wakefulness and decrease during sleep.^44, 45^ In line with this, ECHS1D mice spent significantly more time in the active wake period and reduced time in slow wave sleep. This suggests ECHS1D mice may have sleep disturbances where slow wave power is increased during wake periods. Sleep-disordered breathing is common in Leigh syndrome, although this has not been reported in ECHS1D patients to date.^46, 47^

In addition to sleep disturbances, delta slowing is an indicator of interictal activity.^48, 49^ In agreement with this, ECHS1D mice presented with significantly increased epileptiform discharges and were highly susceptible to seizures provoked by low doses of the chemiconvulsant, PTZ. Many mechanisms have been proposed to explain the relationship between mitochondrial dysfunction and seizure activity, but the exact underpinnings remain undefined. In our model, we observed a trending decrease in astrocytes within aged ECHS1D mice. Astrocytes regulate glutamate uptake and release,^50^ so reduction of astrocytes following ECHS1 loss may contribute to the epileptic phenotype. Subsequent studies are needed to determine if reduced GFAP expression is driven by reduced astrocyte proliferation during development or by increased cell death throughout aging, as both mechanisms are plausible. Increased microglial activation is both a driver and consequence of epilepsy,^51^ and the increase in IbaI staining in ECHS1D mice suggests microglial activation is associated with the epileptic phenotype in this model as well.

ECHS1D mice had exercise intolerance after repeat trials in rotarod or following treadmill endurance. Movement and muscle tone impairments are reported in almost all ECHS1D patients,^3^ and patients with a milder phenotype will report exercise-induced dystonic attacks.^52, 53^ Exercise intolerance in mitochondrial disease patients is primarily driven by impaired respiration capacity within skeletal muscle,^54^ although the central nervous system is also responsible for responding to demands imposed by aerobic exercise.^55^ Recent studies report decreased mitochondrial respiration in patient fibroblasts or ECHS1D cell models.^15–17^ When we assessed ECHS1D mouse liver, we found that in contrast to findings in cell models, OxPhos complex assembly and energy status was normal. We also assessed gluconeogenesis in ECHS1D mice as mitochondrial metabolism regulates liver gluconeogenesis, with altered mitochondrial function increasing or decreasing rates of glucose production.^56, 57^. In ECHS1D mice circulating glucose levels were unchanged between groups, supporting normal gluconeogenesis. Together, these findings suggest that in the absence of stressors, mitochondrial health was largely normal in ECHS1D mouse liver, however, studies are needed to assess mitochondrial function in brain and skeletal muscle.

In testing disease mechanisms, we found a striking worsening of seizure susceptibility in valine treated ECHS1D mice. Patient reports led us to hypothesize that valine, but not leucine or isoleucine, supplement would worsen disease; future work may test this by treating mice with leucine or isoleucine supplement. These results support the hypothesis that impaired valine metabolism is a major driver of pathogenesis, although the variable success in valine restriction in patients suggests this is not the sole mechanism.^10–14^ It remains to be tested whether valine restriction in ECHS1D mice would improve their baseline phenotypes.

ECHS1D patients report symptom onset or worsening following infection^5, 58^ and inflammation drives disease onset in animal models of Leigh syndrome.^35, 59^ In support of the hypothesis that ECHS1 loss results in dysregulated inflammatory signaling, amino acid content was significantly increased in the liver of ECHS1D mice. In rats, sepsis increases liver amino acid uptake and systemic inflammation in humans is associated with increased circulating amino acids caused by elevated skeletal muscle catabolism.^60, 61^ Future studies will quantify amino acid content within muscle and blood to test this. Further, ECHS1D mice displayed sensitivity to inflammatory insults, shown by reduced survival, failure to thrive, and increased seizure susceptibility following LPS treatment. This study was limited by only LPS-treated mice being tested, so future studies will directly compare treated and non-treated mice to validate these findings.

In conclusion, we developed a novel model of ECHS1 Deficiency that displays patient-relevant phenotypes and identified novel ones that may be relevant to human disease. We demonstrate the use of ECHS1D mice in testing disease mechanisms, which can be used in developing therapies for patients that currently have limited treatment options. These findings are directly relevant to ECHS1D and may be applicable to other mitochondrial disorders.

## Data availability

Data that support the findings of this study are available from the corresponding author, upon reasonable request.

## Acknowledgements

We recognize assistance of the Jackson Laboratory Genome Technologies Services in the generation of mice. The Jackson Laboratory scientific services are supported in part through the National Cancer Institute’s Cancer Core Grant P30CA034196. We thank the UT Southwestern’s Histopathology Core for histological services and the UT Southwestern Whole Brain Microscopy Facility (RRID:SCR_017949) for assistance with slide scanning. We thank the UT Southwestern Preclinical MRI Research Core that is partially funded by Cancer Prevention and Research Institute of Texas (RP210099) for MRS analysis. Behavior testing was performed by the staff of the Rodent Behavior Core which is supported by the UT Southwestern Peter O’Donnell Jr. Brain Institute. Endurance testing was performed by the UT Southwestern Metabolic Phenotyping Core which is supported by P01DK119130 and P30DK127984. We thank the UT Southwestern Neuro-models Facility for assistance with telemetry implant surgery and EEG recording (RRID:SCR_022529). Targeted Metabolomics were performed with assistance from Monika Mizerska and Morgan Villega.

## Supplementary material

Supplementary material is available online.

