## Supplemental Data for "Valine and Inflammation Drive Epilepsy in a Mouse Model of ECHS1 Deficiency"

**Supplemental Table 1: Sequences for Mouse Model Generation**

| Oligo | Sequence (5' - 3') |
|---|---|
| sgRNA | TGTACTGAAAGTTAGCACCT |
| Mutagenic Donor[a] | GCGATGTGGTCAGGTTGATGACCAAGATTAACCAACCACGGTGACTTTAATATCTTTATGATGAC ACTTCCCTTTGTCCTCTTGACCTAGGTGC**A**AACT**C**TCAGTA**T**ATCATCACAGAAAAGAAAGG |
| Target Amplification (Fwd) | CTTCCACAGGTAAGGCTTGG |
| Target Amplification (Rev) | CCTTATCCCCACCAGTGAGC |

[a]DNA sequence of the donor oligonucleotide was altered to prevent guide reassociation with edited genomic DNA via the incorporation of three nucleotide changes (identified in bolded font, underlined). Founder mice only contain A31A (c.93T>A) and F33S (c.98T>C) edited alleles.

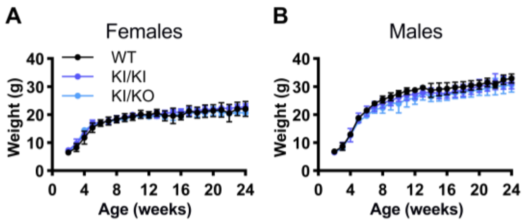


**Supplemental Figure 1: WT and ECHS1D mice have similar body weights**. Mice were weighed weekly beginning at 2 weeks of age and until at 24 weeks of age. There were no differences in weight gain or maintenance between female (A) and male (B) WT and ECHS1D mice.

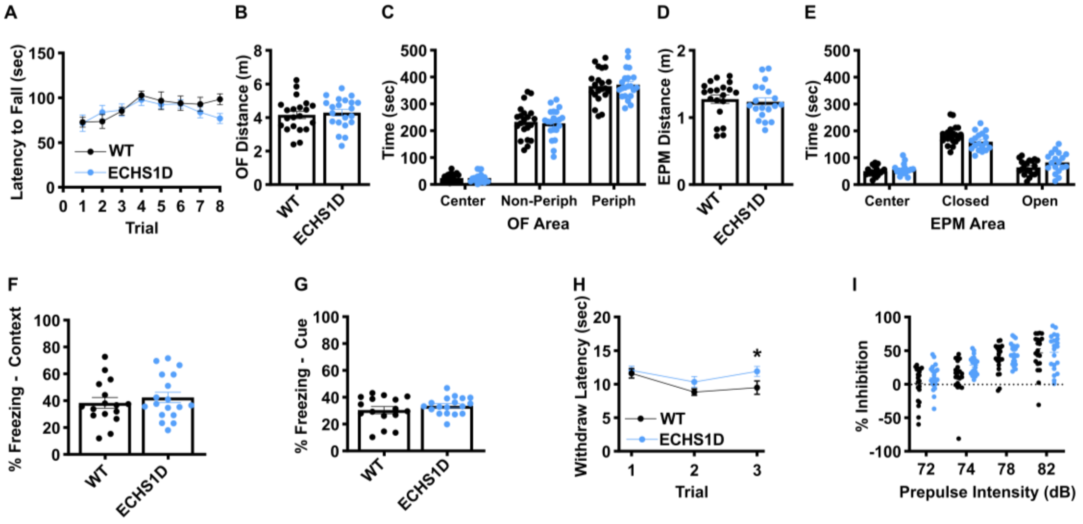


**Supplemental Figure 2: Behavior of ECHS1D mice at 3 months of age.** Beginning at 3 months of age, mice were subjected to a battery of behavioral testing (N=20/genotype). (**A**) Rotarod testing was performed and latency to fall across 8 trials was recorded. (**B-C**) Open field total distance traveled (B) and time spent in each area of the testing arena (C). (**D-E**) Elevated plus maze total distance traveled (D) and time spent in each area of the maze (E). (**F-G**) Fear conditioning percent time spent freezing during the context (F) or cue (G) portion of the test. (**H**) During hotplate testing, the latency to withdraw a paw from the hot surface was recorded. (**I**) Startle responses following an auditory stimulus at 72, 74, 78, or 82 dB was recorded, and percent inhibition was calculated. Each dot represents an individual mouse with SEM indicated. H: Repeat measure two-way ANOVA with Sidak multiple comparisons, WT vs. ECHS1D *$p<0.05$.

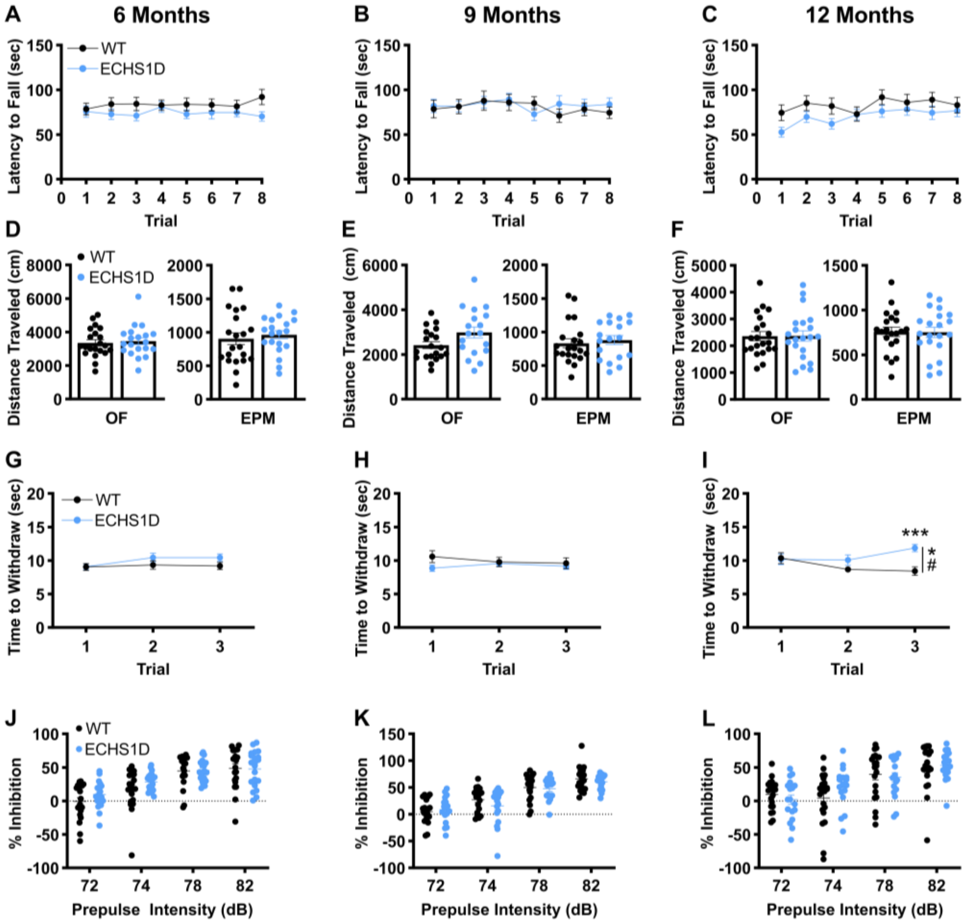


**Supplemental Figure 3: Longitudinal behavioral testing in ECHS1D mice at 6, 9, and 12 months.** Following testing at 3 months of age, WT and ECHS1D mice were subjected to repeat behavioral testing at 6, 9, and 12 months of age (N=20/genotype). **(A-C)** Rotarod testing was performed and latency to fall across 8 trials was recorded. **(D-F)** Total distance traveled in open field and elevated plus maze. **(G-I)** Hotplate testing was performed and latency to paw withdraw from hot surface was recorded across 3 trials. **(J-L)** Percent inhibition was calculated across 4 different prepulse intensities, and there were no differences in inhibition response at any age tested. Each dot represents an individual mouse with SEM indicated. I: Repeat measure two-way ANOVA with Sidak multiple comparisons, Trial x Genotype, *$p<0.05$, Genotype effect, $^{\#}p<0.05$; WT vs. ECHS1D ***$p<0.001$.

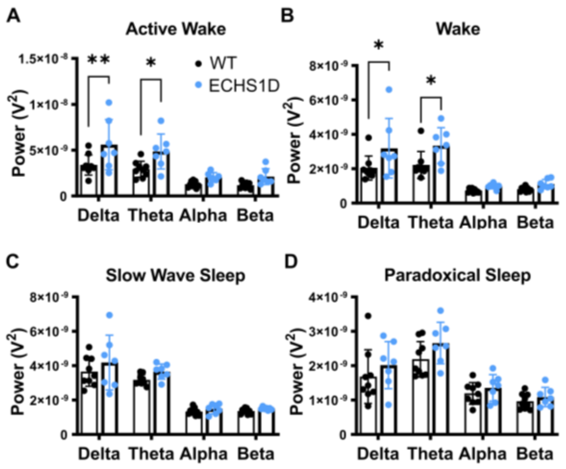


**Supplemental Figure 4: EEG power across different sleep states.** At ~3.5 months of age, mice received wireless telemetry implants and EEG activity was recorded over a 24-hour period (N=7-9/genotype). (**A-D**) Sleep staging was performed and total EEG power from delta, theta, alpha, or beta frequencies was calculated within each sleep stage including active awake (A), wake (B), slow wave sleep (C), and paradoxical (D). Each dot represents an individual mouse with SEM indicated. Two-way ANOVA with Sidak multiple comparisons, *p<0.05, **p<0.01.

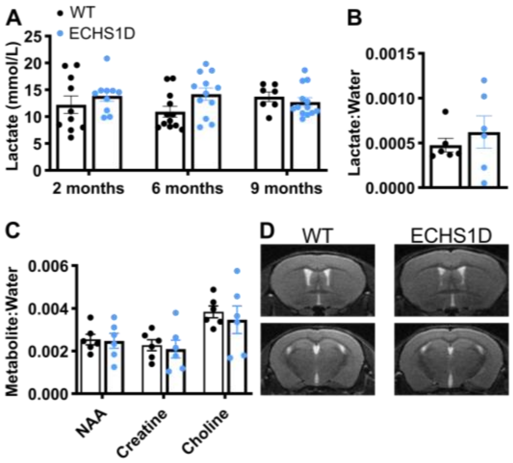


**Supplemental Figure 5: Serum and brain lactate levels in ECHS1D mice.** (**A**) Lactate was quantified from serum isolated at 2, 6, and 9 months of age from WT or ECHS1D mice. **(B-D)** At 9-months of age, WT and ECHS1D mice were subjected to MRS quantification of brain lactate (B), N-acetylaspartate (NAA), creatine, and choline (C). Metabolites were normalized to the signal from water. Each dot represents an individual mouse with SEM indicated.

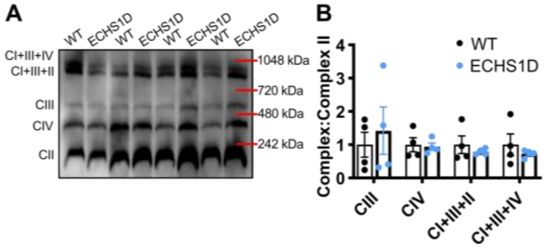


**Supplemental Figure 6: Mitochondrial OxPhos assembly is normal in ECHS1D mice.** (**A-B**) Mitochondria was isolated from the liver of 12-month-old WT and ECHS1D mice and OxPhos complex assembly was examined on a native gel (A) and quantified (B) (N=4/genotype). There were no differences in individual complex or supercomplex assembly.

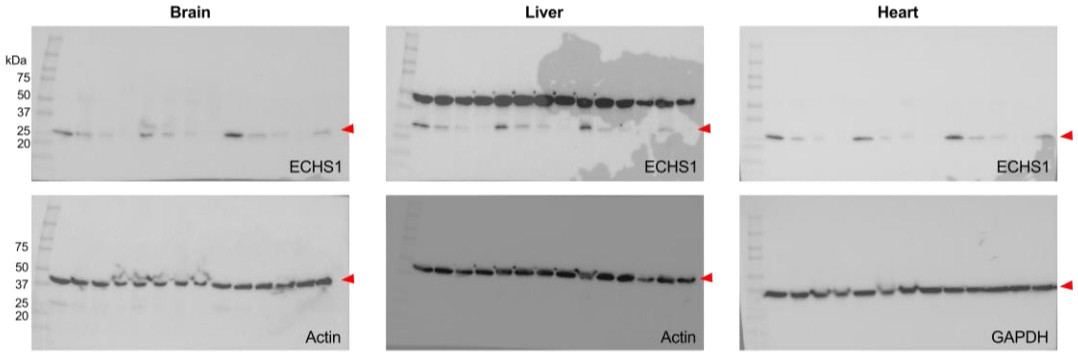


**Supplemental Figure 7: Uncropped blots use in Figure 1.** Full blots that were cropped for clarity in Figure 1. Both ECHS1 and housekeeping proteins were probed on the same blot for each tissue. Red arrows indicate the band of interest.
